# Direct Binding of miR-155 to FLT3 Regulates Key Cellular Functions in Acute Myeloid Leukemia

**DOI:** 10.64898/2026.01.15.696285

**Authors:** Antony Truong, Sarah Warsi, Omeyme Naqchi, Amr Al-Haidari

**Author notes:** Correspondence to: Amr Al-Haidari, PhD, Current affiliation: Department of Urology, University of Rochester Medical Center, Rochester, NY, USA. Independent researcher (Omeyme Naqchi).

## Abstract

Mutation in FLT3 protein is one of the most common mutations in acute myeloid leukemia (AML). Most patients with FLT3-ITD mutation detected at diagnosis or acquired during treatment display poor prognosis and resistance to tyrosine kinase inhibitors or chemotherapy. Existing clinical and pre-clinical data implicate miR-155 in the carcinogenesis of hematological cancers, including FLT3-assocaited AML. However, the role of miR-155 in regulating FLT3-ITD mutation remains elusive. In this study, we have applied loss-of-function studies using wild-type and mutated leukemic cell line models to validate the functional effect of miR-155 inhibition in leukemic cells. Our bioinformatics analysis indicates that FLT3 has a binding site for miR-155 which makes it a direct target of miR-155. Specific targeting of miR-155 by miR-155 inhibitor induced cell apoptosis and reduced FLT3-ITD-mediated cell proliferation and survival. Our data suggests that miR-155 could be a potential therapeutic target for FLT3-associated AML.

## Introduction

The regulation of hematopoietic cell proliferation, survival, and differentiation is tightly regulated by Fms-like tyrosine kinase 3 (FLT3) ^1^. Mutations or over expression of FLT3 gene have been associated with poor prognosis in patients with AML and Acute Lymphoblastic Leukemia (ALL) respectively ^2–4^. The most common FLT3 mutations with clinical relevance to AML are the internal tandem duplication (ITD) mutations of the juxta membrane domain and the tyrosine kinase domain (TKD) of the FLT3 gene ^5^. Of importance, ITD mutations induce constitutive ligand-independent activation of FLT3 which activate multiple signaling pathways, resulting in uncontrolled cell proliferation and up-regulation of anti-apoptotic oncogenes ^6^.

MicroRNAs (miRNAs) are short run of non-coding nucleotides (21–22 nucleotides) that control post-transcriptional gene regulation. It is becoming clear that miRNAs play an instrumental role in cancer progression and metastasis by exerting global genetic and epigenetic changes. In the limelight of miRNAs, miRNA-155-5p (hereafter referred to as miR-155) has been shown to be overexpressed in acute leukemias and play a crucial role in leukemogenesis, leukemia progression, bone marrow niche transformation, and therefore considered as a key player in drug resistance and relapse ^7,8^. One study showed reduced survival and increased apoptosis after treatment with a miR-155 inhibitor in FLT3 mutated AML cell lines in vitro ^9^. Previous reports have also shown that miR-155 alters the function of essential proteins with an established roles in inflammation and malignancy such as RhoA, PAD4, TP53, BRCA1 in sepsis and colon cancer, and in hematopoiesis such as RUNX1 and MLL ^10–14^. The potential clinical relevance of miR-155 has been previously suggested. A systematic review and meta-analysis study conducted by Lu tang et al. found that miR155 levels are associated with unfavorable outcomes and were significantly higher in AML patients compared to healthy controls ^8^. Interestingly, one study demonstrated that patients with myelodysplastic syndrome MDS, a pre-leukemic stage, exhibited lower levels of miR-155 at diagnosis and progressively increased as AML developed ^15^. The high expression of miR-155 was found to be correlated with poor prognosis across multiple adult and pediatric AML cohorts^16–18^. In a study conducted multivariable analyses from patients with cytogenetically normal AML found that miR-155 expression levels were associated with a lower complete remission rate and shorter disease-free survival^19^. In AML, miR-155 has been implicated in various signaling pathways including those with established roles in leukemogenesis such as NF-kB and STAT5 pathways which are involved in FLT3-ITD signaling^20,21^. In this context, pro-proliferative capabilities of FLT3-ITD AML cells were found to have higher levels of phosphorylated AKT which are correlated with reduced miR-155-mediated repression of the hematopoietic transcription factor PU.1, PTEN, and SHIP1^9,20,22^.

Moreover, a previous report demonstrated that FLT3-ITD expression activates STAT5 and RAS/MAPK signaling pathways in stably transfected cell lines as well as in primary AML cells harboring FLT3-ITD mutations^23^. One study conducted by Lingyan Wang et al. showed that miR-155 increased treatment sensitivity of FLT3-ITD AML cells to both chemotherapy and FLT3 inhibitors by targeting PIK3R1, resulting in glycolysis pathway blockage^24^. While these data indicate that miR-155 and FLT-ITD contribute to mutual functional outcomes, the exact relationship between them is still poorly understood. In our study, we show that FLT3-ITD could be a direct target for miR-155 and suggest a therapeutic potential of miR-155 inhibition for leukemic cells harboring FLT3-ITD mutation.

## Materials and methods

### Cell culture

Human MV4-11, Molm-13, and THP-1 leukemia cell lines were cultured in Roswell Park Memorial Institute, RPMI, (VWR Chemical, France) supplemented with 2mM glutamine, 100U/ml Penicillin, and 0.1 mg/ml streptomycin, 10 mM HEPES, and 10% fetal bovine serum (Gibco; Thermo Fisher Scientific, Inc.). All cells were maintained in a humidified atmosphere containing 5% CO_2_ and 37°C.

### Bioinformatics analysis of miR-155 binding site

The 3’-UTR of FLT3 mRNA sequence was retrieved from UCSC Genome Browser on Human (GRCh38/hg38) (Human Hg38 Chr2:25160915-25168903 UCSC Genome Browser V457, n.d.). Manual *in silico* sequence analysis was used to investigate a complimentary based sequence alignment to the seeding region on miR-155. Alignment of at least 7-mer was selected for further investigation as a potential binding site. For multiple sequence analysis, the sequence of miR-155 of different species was retrieved from miRBase (microRNA database) https://www.mirbase.org/ and Clustal Omega pairwise sequence alignment tool of the EMBL-EBI, UK (https://www.ebi.ac.uk/jdispatcher/msa/clustalo) was used for multiple sequence alignment from different species.

### Knockdown of miR-155

MV4-11 cells were seeded in 6-well plate until seeding density reached 60-80% and transfected with 50 nM human miR-155-5p inhibitor (hereafter referred to as AntimiR-155) and AntimiR-155 negative inhibitor control (hereafter referred to Anti-miR155 N.Ctrl) oligonucleotides using RNAimax lipofectamine transection reagent (Thermo Fisher Scientific, Inc.), following the manufacturer’s instructions, and incubated at 37°C with 5% CO_2_ for 24h.

### Trypan blue exclusion assay

A Trypan blue exclusion assay was performed to determine cell viability and to rule out any toxicity caused by the transection reagent after every cell transfection experiment. Cells were stained using 0.4% trypan blue dye and counted using Bürker counting chambers and the viability index (VI) was calculated as the ratio of the number of live cells on wells containing antimiR-155 divided by the number of cells in the control wells.

### Reverse transcriptase Real-Time Quantitative Polymerase Chain Reaction (RT-PCR)

Total RNA was extracted from transfected MV4-11 cells using Quick-DNA/RNA™ Miniprep Plus extraction kit (Zymo Research, Irvine, CA, USA) following the manufacturers protocol. The RNA concentration used was normalized to 60 ng/μl across the samples. For first-strand cDNA synthesis of miR-155, small RNAs were polyadenylated and reverse transcribed using the Mir-X miRNA First-Strand Synthesis kit (Takara Bio inc, USA). For quantitative PCR (qPCR), the concentration of cDNA was standardized to 10 ng/μl and miR155-5p and FLT3 gene expression was performed using SYBR Green qPCR Master Mix (MCE, USA), according to the manufacturer’s instructions. The primers used in the study were previously described and listed in table 1. GPDH and U6 snRNA were used as housekeeping genes, and the relative expression expressed by Fold-change was calculated using the 2^-ΔΔCT^ method.

**Table 1:**
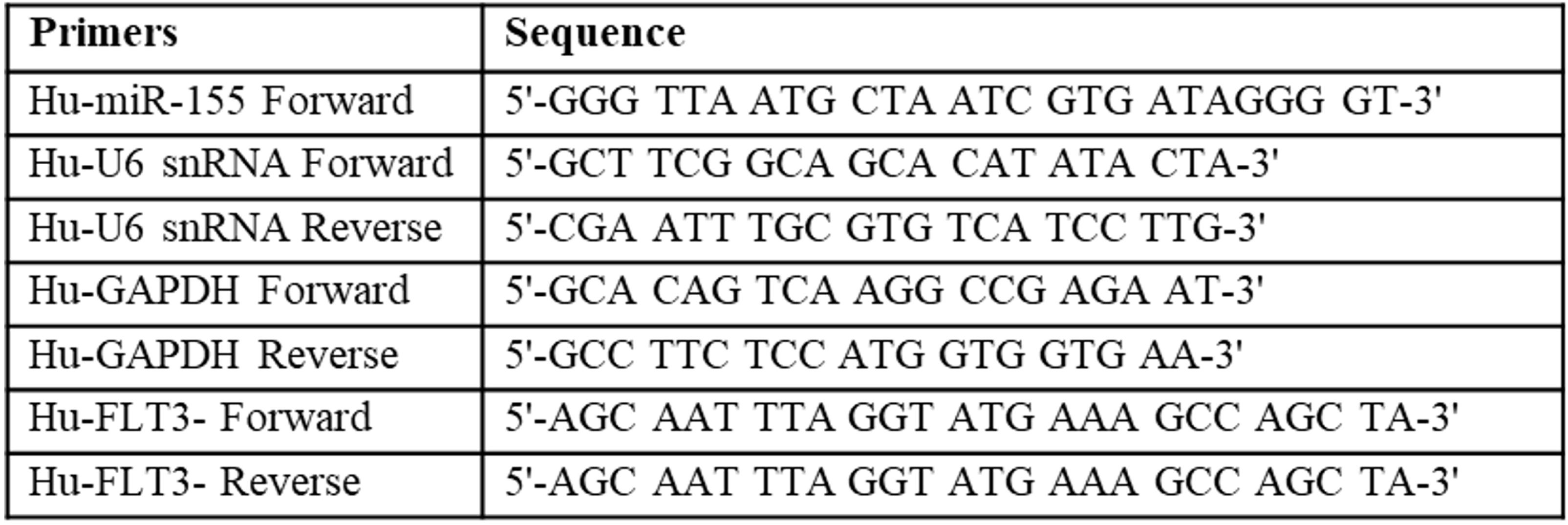
Primers used in qPCR gene expression.

### Protein expression

FLT3 (CD135) expression was evaluated in MV4-11 cells using flow cytometry with or without transfection and treated according to cell transfection described above. After 24h, one million cells were first washed twice with PBS and then resuspended in 1 ml PBS. One hundred μl is then used in triplicates for each sample. Surface staining was achieved by staining cells with 5 μl of PE Mouse Anti-Human CD135 (BD biosciences, Sweden) for 10 minutes in the dark at R.T. For intracellular staining, cells were fixed and permeabilized using FIX & PERM™ cell permeabilization kit (Invitrogen™) according to the manufacturer’s instructions. The cells were then washed twice with 2 ml PBS at 500 xg for 5 minutes. The PBS is then discarded, and the cells were resuspended in 400 μl PBS before measuring CD135 expression using Cytoflex (Beckman coulter, USA) with 10,000 events being recorded on fast flow. Unstained cells were used as a negative control, and data were analyzed using FCS express (De Novo Software v7.0, USA).

### Functional studies

All functional studies underwent treatment of antimiR-155 according to the transfection protocol described above. The effect of antimiR-155 knockdown on MV4-11 cells was demonstrated using Apoptosis, proliferation, and colony formation assay kits as follows:

#### 1. Cell proliferation assay

Cell proliferation was evaluated in quadruplicates using Cell Counting Kit-8 assay (CCK-8; BosterBio). Briefly, cells were transfected with 100 nM antimiR-155 or antimiR-155 N.Ctrl as described above for 24h, and then 5000 cells were seeded in 96-well culture plate and incubated for 24h at 37 °C and 5 % CO_2_. To quantify proliferation, 10 μL of CCK-8 was added per well for 2 h and absorbance at 450 nm was measured using GloMax® Discover Microplate Reader (Promega Corp). Proliferation % was calculated by dividing the OD absorbance of cells in wells containing antimiR-155by the OD of cells in the control wells.

#### 2. Detection of apoptosis

The quantification of cell death was determined using annexin V-based flow cytometry assay. Cells were transfected for 24h with AntimiR-155 and AntimiR-155 N.Ctrl as described above and apoptosis was evaluated using CoraLite® Plus 488-Annexin V and PI Apoptosis Kit (Proteintech, IL, USA). Suspension of 1×10^6^ cells/ml was prepared and 100 µl in triplicates of each sample was dispensed to FACS tubes. CoraLite Plus 488-Annexin V and PI were used to stain the cells for 10 min in dart at R.T. Cells were then washed with PBS and resuspended in 400 µl PBS and apoptosis was measured using Cytoflex flow cytometry (Beckman coulter, USA) with 10,000 events being recorded on fast flow. Unstained cells were used as a negative control, and data were analyzed using FCS express (De Novo Software v7.0, USA).

#### 3. Colony Formation assay

The medium methylcellulose was prepared according to MethoCult™ H4100 protocol. One thousand MV4-11 cells were seeded/well on a 6-well plate after they had been transfected with antimiR-155 or antimir-155 N.C for 24h. Cells were seeded in a ratio of 1:10 cells to methylcellulose media. Briefly, 400μl of transfected cells containing 1x10^5^ cells/ml were collected to methylcellulose media in a final volume of 4ml and 10%FBS. The samples were then vortexed and left to stand for 5 minutes to settle any air bubbles and then incubated for 7 days at 37°C, 5% CO_2_. Visible colonies were counted macroscopically and expressed as a number of colonies produced by the cells.

### Data analysis

All data are presented as mean ± standard error of the mean (SEM). Comparison between the treated and control treated groups was achieved using the Mann Whitney nonparametric test. The statistical analysis was performed using Graph prism8 software and P value <0.05 was considered statistical significance.

## Results

### MiR-155 is overexpressed in FLT3-mutated AML cell lines

The overexpression of miR-155-5 has been well established previously in patients with AML^7,8^. However, no evidence whether miR-155 overexpression is associated with increased expression of FLT3 as a result of direct binding and regulation. To address this question, we have used three cell lines where the FLT3 gene has either 2 alleles ITD mutated (MV4-11 cells) or one allele (Molm-13 cells) or wild type; both alleles are normal with no ITD mutation (THP-1 cells). In line with the existing clinical reports, our study showed that miR-155 and FLT3 mRNA are overexpressed in FLT3-mutated cells, MV4-11 and Molm-13 cell lines, compared to wild-type FLT3 cells, THP-1 cells (Figure 1 A &B). In addition, FLT3 protein showed higher expression in FLT3-mutated cells compared to FLT3 wild-type cells, suggesting possible correlation between FLT3 and miR-155 (Figure 1C &D).

**Figure 1.**
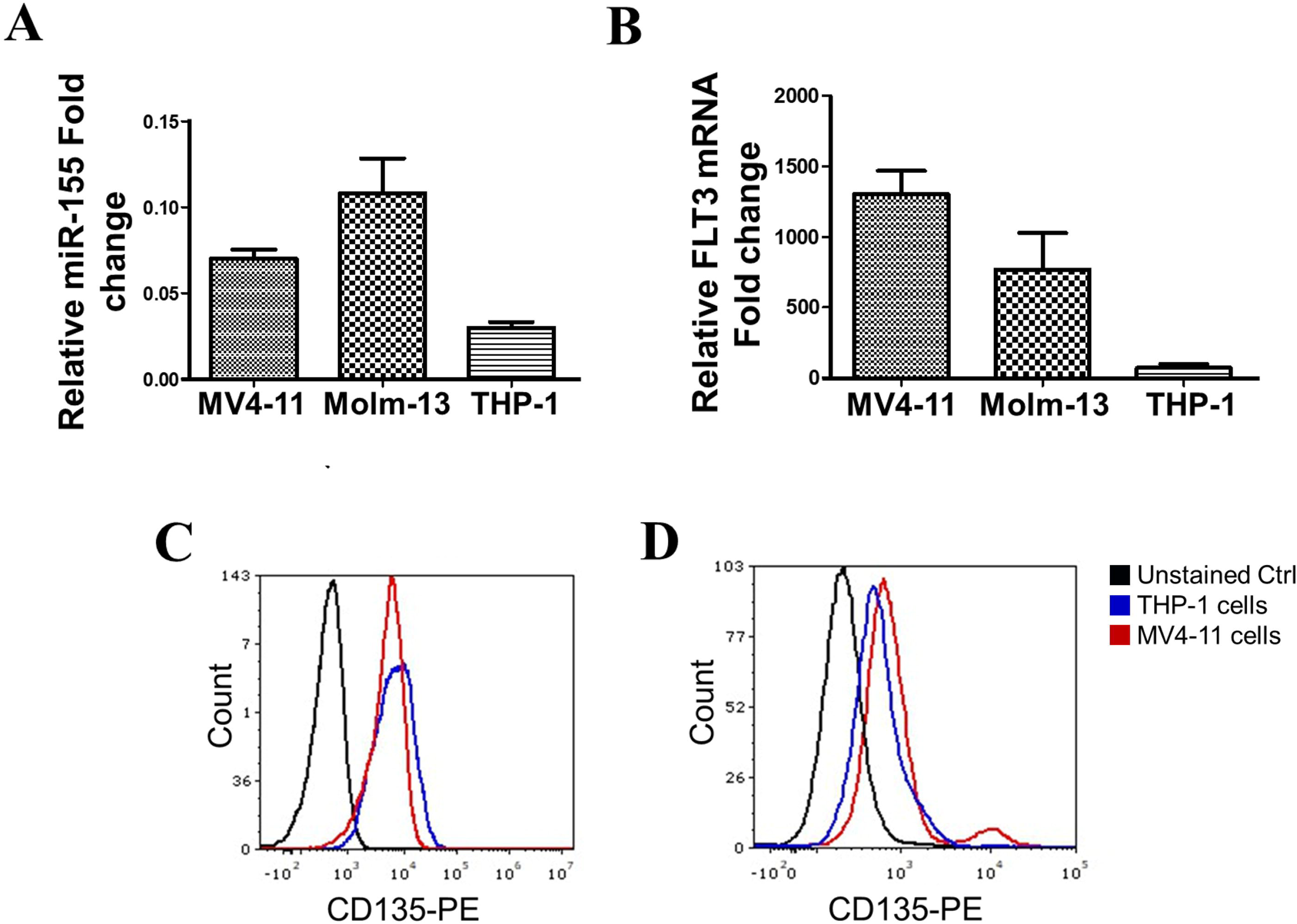
Expression of miR155 and FLT3 mRNA in leukemic cell lines. **A.** miR-155 expression levels were quantified in MV4-11, Molm-13, THP-1 cell lines by qPCR, and U6 expression was used as endogenous control. **B.** mRNA expression levels of FLT3 in MV4-11, Molm-13, THP-1 cell lines. The GAPDH gene was used as an endogenous control. **C.** Surface and **D.** intracellular expression of CD135 (FLT3) in MV4-11 and THP-1 cells. Cells were stained with anti-human CD135-PE, and unstained control was used to rule out any background noise posed by cell autofluorescence. Data are presented as Mean ±SEM of 3 independent experiments, and qPCR data represented as fold-change.

### FLT3 is a target of miR-155

Given the possible correlative data between FLT3 and miR-155 expression, we sought to establish the association between miR-155 and FLT3 and whether FLT3 is a direct target of miR-155. Thus, we used complementary-based nucleotide bioinformatics analysis to examine the 3’-UTR region of FLT3 mRNA and the seeding region of miR-155. Our bioinformatics analysis showed that FLT3 mRNA contains a putative binding site that is complementary to miR-155 7-mer seeding region, suggesting possible direct binding of miR-155 to FLT3 mRNA (Figure 2A). We have further analyzed this region across different species where we observed that the seeding region of miR-155 is evolutionary conserved across different species, indicating that miR-155 seeding region is crucial in regulating bound mRNAs (Figure 2B).

**Figure. 2.**
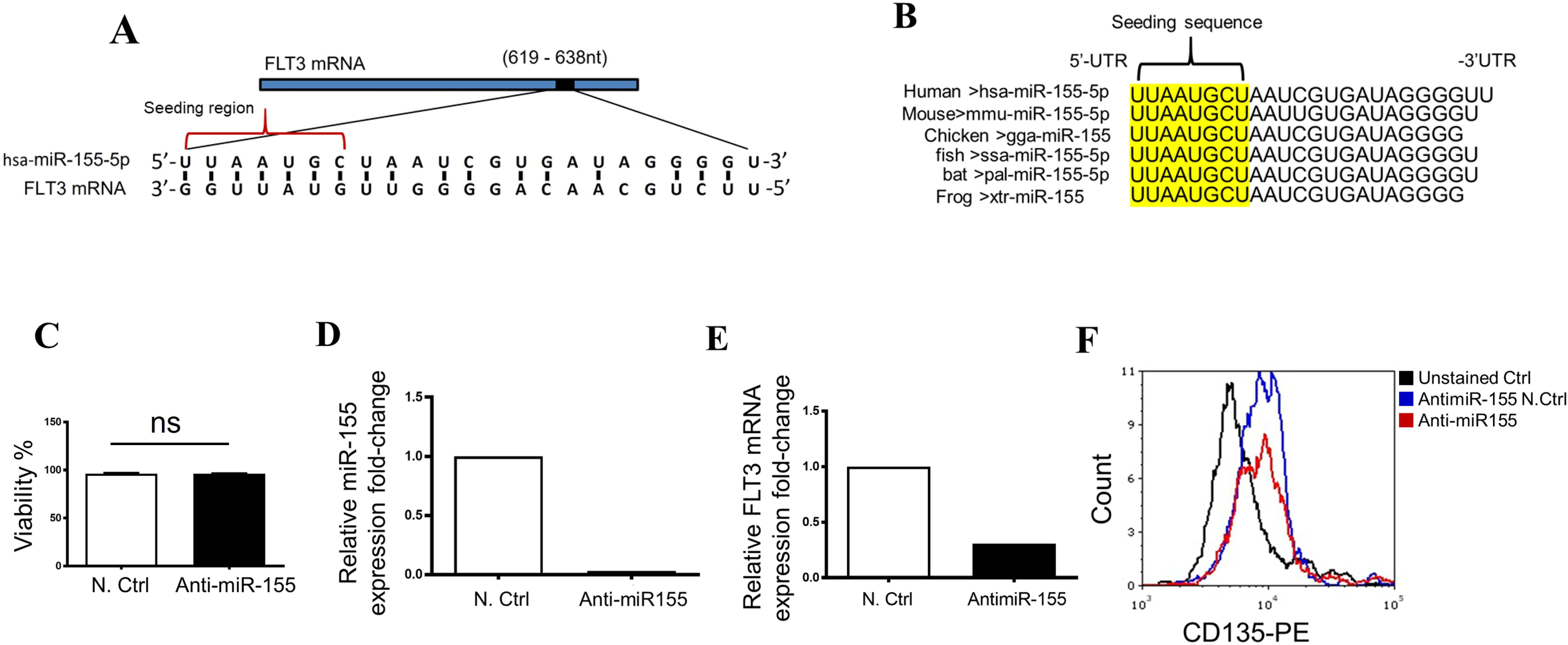
FLT3 is a direct target of miR-155. **A.** Bioinformatics analysis predicted the presence of a putative binding site in FLT3 mRNA 3’-UTR. *In silico* sequence complementary pairwise analysis with 7-mer match between miR-155 seeding region and FLT3 mRNA. **B.** Multiple sequence alignment of miR-155 seeding region across different species. **C.** Cell viability detected using trypan blue exclusion assay after 24h of miR-155 knockdown in MV4-11 cells using 50 nM antimR-155 or antimiR-155 control (Ctrl). **D.** miR-155 expression levels by qPCR in MV4-11 cells 24h post-treatment with 50 nM antimiR-155 or antimiR-155 Ctrl. **E.** FLT3 mRNA expression levels in MV4-11 cells 24h post-treatment with 50 nM antimiR-155 or antimiR-155 Ctrl. The GAPDH and U6 genes were used as endogenous controls in FLT3 and miR-155 qPCR respectively, and qPCR data represented as fold-change. **F.** FLT3 protein expression 24h post-treatment with 50 nM antimiR-155 (red) or antimiR-155 Ctrl (blue). Live cells were stained with anti-human CD135-PE (FLT3) and analyzed using flowcytometry.

### Knocking down miR-155

To address the regulatory function as well as the correlation between miR-155 and mutated FLT3 mRNA, we have knocked down miR-155 using miR-155 specific inhibitor in MV4-11 cells using lipofectamine transfection technology. Our transfection process showed that nearly all cells were intact during the transfection process, indicating that the transfection was not toxic to the cells (Figure 2C). To verify miR-155 knocking down, we used qPCR to examine the expression of miR-155 after transfection with anti-miR-155 (10nM and 50nM) after 24h (Figure S1). The data showed that miR-155 inhibitor 50nM almost abolished miR-155 (Figure 2D). The effect of miR-155 knockdown was further verified on the protein and gene levels of FLT3. We found that knocking down miR-155 reduced FLT3 mRNA by >2-folds and decreased FLT3 protein expression (Figure 2E &F), suggesting a causal relationship between miR-155 and FLT3.

### MiR-155 regulates FLT3-dependent AML cell functions

FLT3 protein regulates essential cell functions such as proliferation and cell survival. To confirm the positive correlation effect between miR-155 and FLT3, we knocked down miR-155 and examined FLT3-dependent cell functions in MV4-11 cell line. Inhibiting miR-155 significantly reduced FLT3-ITD-dependent cell proliferation and decreased the number of colonies as well as induced cell apoptosis (Figure 3A, B &C). This data indicates that miR-155 might be involved in direct regulation of FLT3-mediated cell functions.

**Figure. 3.**
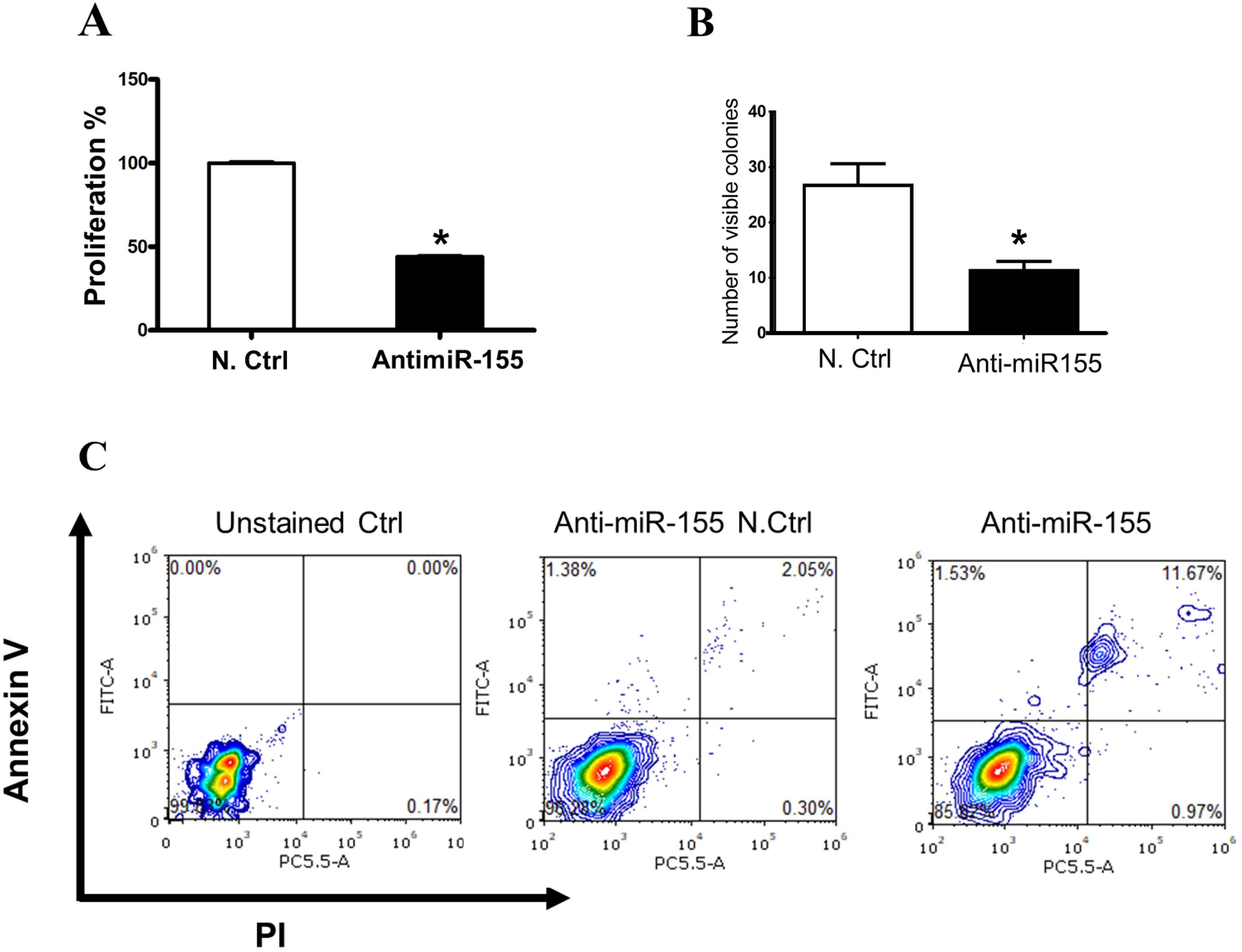
MiR-155 regulates FLT3-ITD-induced cell proliferation and survival. **A.** MV4-11 cell proliferation 24h post-treatment with 50 nM antimiR-155 or antimiR-155 Ctrl. Cells were treated with indicated dose of antimiR-155 or Ctrl and seeded in 96-well plate and incubated 24h. Cell proliferation was then measured using CCK-8 assay. **B.** Colony formation assay was performed after cells were treated with 50 nM antimiR-155 or antimiR-155 Ctrl and seeded in a ratio of 1:10 cells to methylcellulose media and then incubated for 7 days. Visible colonies were counted macroscopically and expressed as a number of colonies produced by the cells. **C.** MV4-11 cell apoptosis was measured by flow cytometry utilizing annexin V and PI markers 24h post-treatment with 50 nM antimiR-155 or antimiR-155 Ctrl. Data represent the Mean ±SEM; significant differences relative to control group are shown as *p<0.05 of three independent experiments.

## Discussion

FLT3-ITD mutation is one of the most detected cytogenetic abnormalities in patients with AML. Patients with positive FLT3-ITD mutation at diagnosis are often associated with poor prognosis and short survival ^3^. Although abnormal high expression of FLT3 in patients with AML has made FLT3 a promising therapeutic target including those tyrosine kinases against FLT3, patients display resistance after treatment and develop relapse with subsequent dramatic progression of the disease. Therefore, new strategies to overcome FLT-3 therapy resistance are greatly needed.

MiRNAs are a family of small non-coding RNAs that control post-transcriptional regulation of gene expression. The role of miRNAs in various diseases has been well documented with prominent dysregulation characteristic features found in many types of cancers ^25^. MiRNAs can act as an oncomiRNAs by targeting oncogenes or tumor supressor-miRNAs by targeting tumor suppressor genes, thus promoting or repressing carcinogenesis process. In AML, miRNA expression landscape suggested that miRNAs might play an important role in the molecular pathogenesis of AML ^26^. Several miRNAs have been implicated in FLT3-ITD^+^ AML including miR-155 ^27–29^. For example, previous reports showed that miR-155 promotes FLT3-ITD–induced myeloproliferative disease through inhibition of the interferon response ^9^. Another study demonstrated that loss of miR-155 sensitizes FLT3-ITD^+^ AML to chemotherapy ^24^. Furthermore, Dennis Gerloff et al. showed that miR-155 mediates NF-κB and STAT5 FLT3-ITD Signaling ^20^. While these reports indicated that FLT3-ITD and miR-155 are involved in AML leukemogenesis, no evidence up to date has demonstrated a direct relationship between miR-155 and FLT3-ITD. In this study, we sought to understand how miR-155 regulates FLT3-ITD-related leukemic cellular functions.

Our study showed that FLT3 mRNA is more expressed in cells with either homozygous or heterozygous mutant allele of FLT3-ITD. For example, we found that FLT3 has higher mRNA expression in MV4-11 (both alleles are mutant), and Molm-13 cells (one allele is mutant) compared to the wild-type FLT3 in THP-1 cells. In addition, miR-155 expression displays higher levels in MV4-11, and Molm-13 cells compared to THP-1 cells. These results indicate that mutated FLT3 and miR-155 might be positively correlated. To address this issue, we have selected MV4-11 cells for subsequent studies to ensure the full penetrant effect of the mutation on the FLT3-ITD/miR-155 axis-mediated cell functional outcomes. FLT3 protein expression was also higher in MV4-11 mutant FLT3 cells compared to THP-1 with wild-type FLT3 cells. Therefore, miR-155 was knocked out using a specific miR-155 inhibitor and subsequent functional analyses were examined. Transfecting cells with miR-155 inhibitor had no impact on cell viability and almost abolished miR-155 expression and reduced FLT3 mRNA by >2 folds. Using flow cytometry, we were able to observe a decrease in FLT3 protein expression under miR-155 inhibition. This suggests that FLT3 and miR-155 have a mutual positive correlation. A large body of clinical evidence demonstrated that higher expression of miR-155 is associated with cytogenetically positive FLT3-ITD AML patients ^29^. Moreover, previous reports showed that knocking out of miR-155 using CRISPR-Cas9 increased sensitivity to FLT3 inhibitors in MV4-11 cells ^24^. One in vivo study used anti-miR155 to block miR-155 in 32D cells which were injected in mice found significant reduction of accumulated leukemic cells in the bone marrow^30^. These data are in line with ours which support the positive correlation between FLT3 and miR-155.

Given the observed positive correlation between FLT3 and miR-155, we have asked a question whether FLT3 is a direct target for miR-155. To further address this relationship, we made a manual seed-sequence complementary-based bioinformatics analysis and found that that the 3’-UTR of the FLT3 mRNA contains a putative binding site that is complementary to the seeding region of miR-155, suggesting that miR-155 binds directly to FLT3 mRNA. Moreover, the seeding region of miR-155 has been found to be evolutionary conserved across different species, indicating the complex role of miR-155 in various biological systems. Notably, this binding is not reported by commonly used miRNA target prediction algorithms. The absence of this binding site from these prediction tools might reflect the inherent limitations of current algorithms, which are likely based on strict mer-based canonical site signature coupled to scoring thresholds and reference transcript annotations. Although the identified miR-155 7-mer binding site in the 3’-UTR of FLT3 is evolutionarily conserved, it may not satisfy all algorithm-specific criteria required for inclusion, such as optimal scores or canonical seed or structural energy classification.

Several studies which highlighted the role of miR-155 on FLT3 only demonstrated the effect mediated through intermediate proteins. In other words, miR-155 might indirectly regulate FLT3. For example, one study demonstrated that miR-155-associated FLT3-ITD expression can directly target PU.1, a transcriptional factor involved in myeloid differentiation, which in turn blocks proliferation and induces apoptosis ^20^. In addition, miR-155 has been also implicated in the regulation of FLT3-ITD via NF-κB, and targeting NF-κB using MLN4924 (Pevonedistat) was associated with reduced miR-155 expression and delayed tumor progression in vivo ^22^. Although these data demonstrated some regulatory role of miR-155, they also reflect the indirect mechanism by which miR-155 mediates FLT3-ITD regulation. In contrast, our data showed that miR-155 can bind FLT3 directly and provide evidence of functional validation. It is well-documented that FLT3-ITD promotes cell proliferation and survival independently of its ligand due to constitutive phosphorylation and activation of FLT3 protein imposed by ITD mutation ^6^. Our data showed that inhibition of miR-155 significantly reduced cell proliferation and colony formation. Although these outcomes are not surprising when targeting FLT3, reduced proliferation and survival mediated by suppression of miR-155 suggest that FLT3 is tightly regulated by miR-155. Moreover, targeting miR-155 by a specific miR-155 inhibitor induced apoptosis in FLT3-ITD^+^ MV4-11 leukemic cells. Our data are in line with literature evidence which showed that inhibition of miR-155 leads to impairment of collagenic capabilities, reduced cell proliferation, and induced cell apoptosis. For example, one study demonstrated that miR-155 increased proliferation of AML and targeting miR-155 resulted in decreased cell proliferation by reducing the growth-inhibitory effects of the interferon (IFN). Further analysis indicated that the inhibitory effect was mediated by direct miR-155 targeting CCAAT/Enhancer-Binding Protein Beta (Cebpb) ^9^. Another study showed that f miR-155 decreased cell proliferation in MV-4-11 and MOLM-13 cells and the effects were associated with increased apoptosis. Additional analysis indicated that miR-155 act on WEE1 protein, a nuclear kinase belonging to the Ser/Thr family of protein kinases that regulate cell cycle ^31^. While these lines of evidence clearly indicate that targeting miR-155 regulates FLT3-ITD-mediated cellular functions, it also explicitly showed that these effects are more likely a result of intermediate protein effect mediated by miR-155 regulation and not associated with direct miR-155 binding and regulation of FLT3-ITD. Although our study shows that miR-155 binds directly to FLT3 and mediates functional regulation in AML cell model of ITD mutation, it is still crucial to experimentally validate the binding site before drawing firm conclusions. Nevertheless, this report provides the first evidence of direct targeting of miR-155 on FLT3 by direct binding and adds a supporting ground to the existing literature for future evaluation as a potential AML targeted therapy.

## Contribution

A. AL conceptualized, supervised, conducted, and wrote the original manuscript draft. A. T conducted the study experiments, reviewed the manuscript; S.W conducted the study experiments, reviewed the manuscript; O. N conducted study experiments and reviewed the manuscript.

## Acknowledgement

The authors would like to thank Associate Professor Julhash U. Kazi, Division of Translational Cancer Research, Lund University, for providing us with the leukemic cell lines. We are also grateful to the clinical hematopathology lab staff at Region Skåne for all kinds of assistance. We also would like to thank all those with direct or indirect contributions to support this study, including Skåne University Hospital, the Tore Nilsons Stiftelse, and The Royal Physiographic Society in Lund.

