## Supplementary figure legend for "Direct Binding of miR-155 to FLT3 Regulates Key Cellular Functions in Acute Myeloid Leukemia"

**Supplementary data**

**Figure S1. Inhibition of miR-155 in MV4-11 leukemic cells.** Cells were transfected with anti-miR155 or anti-miR-155 N. Ctrl (10nM and 50nM) for 24h. Cells were then harvested and relative miR-155 expression was determined using qPCR. Data were expressed as a fold-change relative to the control and n=3 of three independent experiments. U6 was used as a housekeeping gene.
