## Supplementary figures and images for "Direct Binding of miR-155 to FLT3 Regulates Key Cellular Functions in Acute Myeloid Leukemia"

### Figure S1

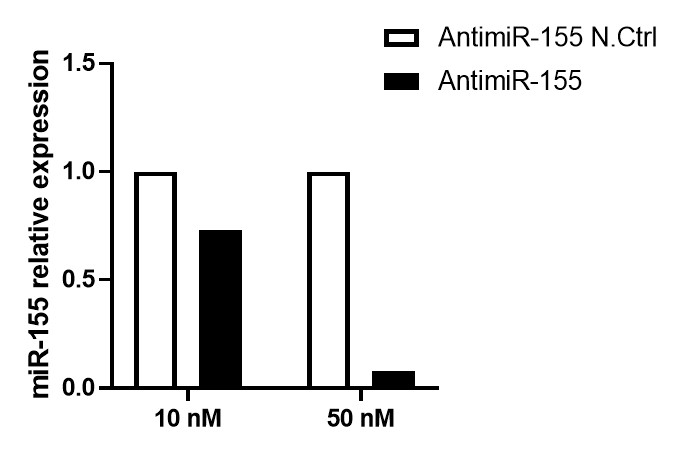
